# The striking *flower-in-flower* phenotype of *Arabidopsis thaliana* Nossen (No-0) is caused by a novel *LEAFY* allele

**DOI:** 10.1101/535120

**Authors:** Anne Mohrholz, Hequan Sun, Nina Glöckner, Sabine Hummel, Üner Kolukisaoglu, Korbinian Schneeberger, Klaus Harter

**Affiliations:** Department of Plant Physiology, Center for Plant Molecular Biology, Universität Tübingen, 72076 Tübingen, Germany; Department of Plant Developmental Biology, Max Planck Institute for Plant Breeding Research, 50829 Cologne, Germany

**Keywords:** *Arabidopsis thaliana*, floral development, flower morphology, *Ds* transposon, classical/sequencing-based mapping, *LEAFY*, DNA-binding

## Abstract

**Summary:** The transition to reproduction is a crucial step in the life cycle of any organism. In *Arabidopsis thaliana* the establishment of reproductive growth can be divided into two phases: In the first phase, cauline leaves with axillary meristems are formed and internode elongation begins. In the second phase, lateral meristems develop into flowers with defined organs. Floral shoots are usually determinate and suppress the development of lateral shoots. Here, we describe a *Ds* transposon insertion mutant in the Nossen (No-0) accession with severe defects in floral development and flower morphology. The most striking aspect is the outgrowth of stems from the axillary bracts of the primary flower carrying terminal secondary flowers. Therefore, we named this mutant *flower-in-flower* (*fif*). However, the insertion of the transposon in the annotated gene is not responsible for the *fif* phenotype. By means of classical and genome sequencing-based mapping, the mutation responsible for the *fif* phenotype was found to be in the *LEAFY* (*LFY*) gene. The mutation, a G-to-A exchange in the second exon of *LFY*, creates a novel *lfy* allele and causes a cysteine-to-tyrosine exchange in the α1-helix of the LFY DNA-binding domain. Whereas subcellular localization and homomerization are not affected, the DNA-binding of LFY^FIF^ is abolished. We propose that the amino acid exchange interferes with the cooperative binding of LFY to its target DNA. To generate the strong *fif* phenotype, LFY^FIF^ may act dominant-negatively by either forming non-binding LFY/LFY^FIF^ heteromers or by titrating out the interaction partners, required for LFY function as transcription factor.

**Significant Statement:** The *fif* phenotype of *Arabidopsis thaliana* No-0 is caused by a novel allele of the *LEAFY* gene

## Introduction

The development of flowers is indispensable for the reproductive success of angiosperm plants. During vegetative growth, the shoot apical meristem (SAM) develops leaves and/or branches, the latter with their own SAMs. After the switch to reproductive growth, the apical meristems give rise to flowers. Floral development differs crucially from vegetative shoot growth, as the flower possesses several types of organs of which the number, arrangement and morphology are species-specific. Furthermore, the development of lateral shoots is inhibited in flowers and floral shoots are determinate after the last reproductive organs have been initiated (Piñeiro and Coupland, 1998; Ma, 1998; Pidkowich *et al.*, 1999). Thus, the coordination of complex molecular processes is necessary for successful floral development. There has been significant progress in recent years towards understanding the molecular mechanisms underlying flower formation. Central to this was the identification and cloning of the genes that initiate and maintain floral development in plant species, including *Arabidopsis thaliana*. The most intriguing discovery was the *Arabidopsis* loss-of-function mutants with structures that are intermediate between floral and vegetative shoots. The cloning of the corresponding genes revealed the existence of the master regulators required for the floral initiation process (FLIP). To date, five FLIP regulatory master genes are known: *LEAFY* (*LFY*), *APETALA1* (*AP1*), *CAULIFLOWER* (*CAL*), *APETALA2* (*AP2*) and *UNUSUAL FLORAL ORGANS* (*UFO*) (Pidkowich *et al.*, 1999). *LFY* and *AP1* play a primary role in initiating the floral program, as the corresponding loss-of-function mutants do not generate shoots with floral characteristics and the ectopic expression of either gene induces precocious flower formation (Irish and Sussex, 1990; Huala and Sussex, 1992; Bowman *et al.*, 1993). Based on its amino acid similarity and expression characteristics *CAL* appears to be functionally redundant to *AP1* (Kempin *et al.*, 1995). *LFY, AP1* and *CAL* encode for transcription factors and are expressed predominantly in floral primordia (Weigel *et al.*, 1992; Mandel *et al.*, 1992; Kempin *et al.*, 1995).

During plant vegetative growth, *LFY* expression increases in newly formed leaves until a certain threshold is reached (Bowmann *et al.*, 1993). LFY then induces the expression of *AP1*/*CAL* genes by binding to the *AP1*/*CAL* promoters. Through their mutual transcriptional up-regulation, LFY and AP1/CAL cooperate to cause the floral transition (Blazquez *et al.*, 2006). Once the floral meristem is established, the FLIP gene functions govern its spatial patterning by inducing the expression of the floral homeotic *ABC* genes, such as *AP2, AP3, Pistillata* (*PI*) and *AGAMOUS* (*AG*). The *ABC* gene functions in turn control the identity of the stereotypically arranged *Arabidopsis* floral organs (Coen and Meyerowitz, 1991; Lohmann and Weigel, 2002). In the course of our study of the influence of abiotic stress on flower symmetry, we searched for novel insertion mutants with defects in floral development or morphology in different *Arabidopsis thaliana* accessions. We focused on genes that had not yet been linked to flowering. A *Ds* transposon insertion mutant, which developed secondary inflorescences with partially aberrant flowers, was identified in the No-0 accession. The wild-type allele of the gene carrying the *Ds* transposon codes for a cystein/histidine-rich C1 domain protein (Shinya *et al.*, 2007; Miwa *et al.*, 2008). However, a thorough genetic analysis revealed that the transposon-inserted allele is not the cause of the observed floral phenotype. Using classical mapping and mapping-by-sequencing, we eventually found a novel mutant allele of *LFY* to be responsible for the aberrant floral development and flower morphology and determined the molecular reason for LFY malfunction.

## Results

### The *flower-in-flower* (*fif*) transposon insertion line displays a novel flower phenotype

In order to identify novel *Arabidopsis thaliana* mutants with defects in flowering we screened the RIKEN Arabidopsis Phenome Information Database (RAPID; Kuromori *et al.*, 2006). RAPID also covers a *Ds* transposon mutant collection in the *Arabidopsis* Nossen-0 (No-0) background (Ito *et al.*, 2002; Kuromori *et al.*, 2004). We identified a transposon-tagged line (15-3794-1), which developed secondary inflorescences with partially aberrant flowers (Fig. 1a). Because of this phenotype, we named this novel *Arabidopsis* mutant *flower-in-flow*er (*fif*).

**Figure 1.**
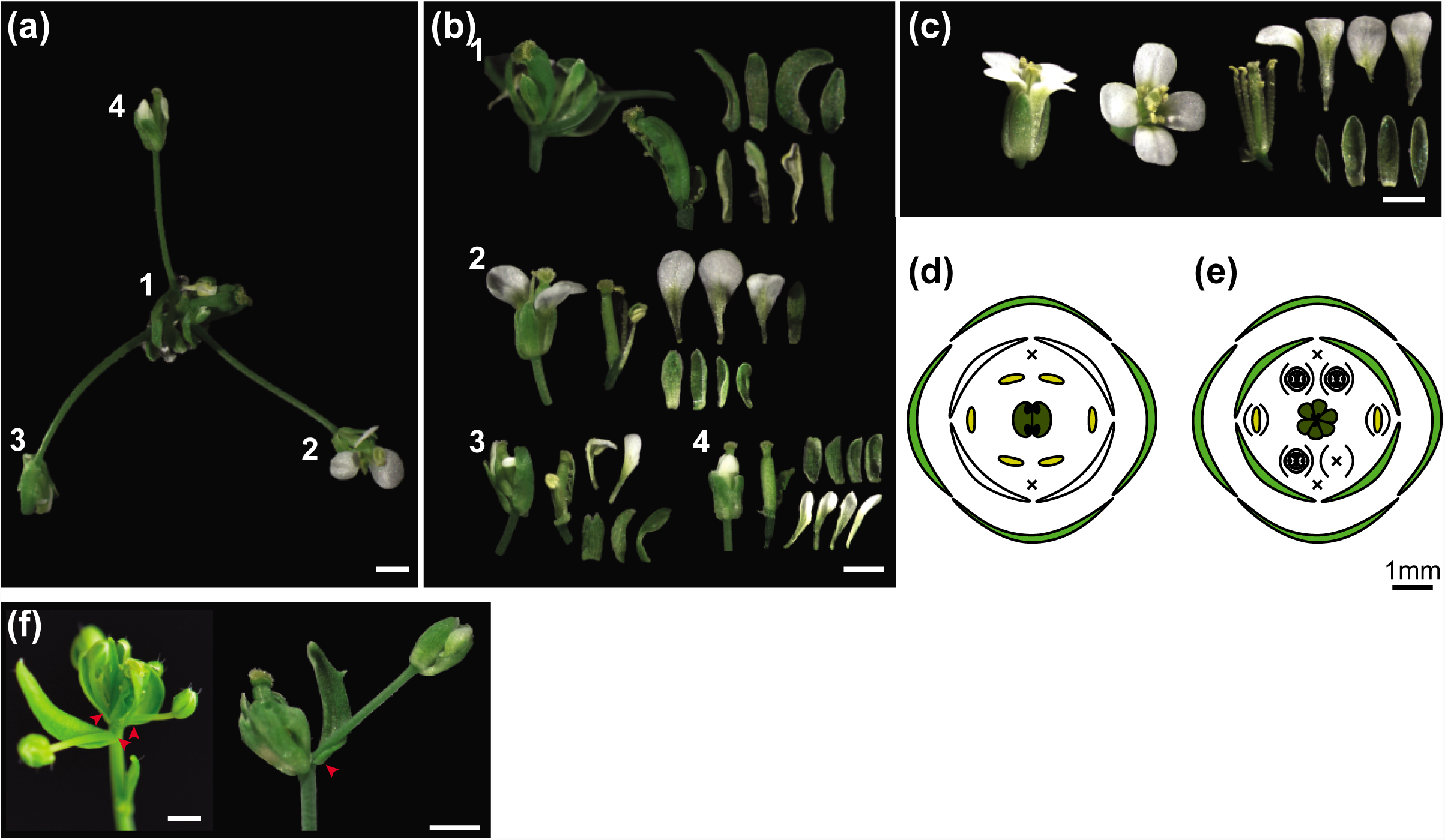
Flower phenotype of the *Arabidopsis thaliana* (No-0) *flower-in-flower* (*fif*) mutant. (a) Overview over representative *fif* mutant “inflorescence” displaying different flower types 1 to 4. (b) Floral organs of the primary *fif* flower (1) and different secondary *fif* flowers (2-4). (c) Flower of the wild-type No-0 accession. (d-e) Flower diagram of the wild-type No-0 flower (d) and the primary flower of the *fif* mutant (e). (f) Primary flowers of the *fif* mutant with stems that outgrow from axillary bract meristems (red arrow heads) and carry secondary flowers. Size bar: 1 mm.

As shown in Figure 1c and d, wild-type *Arabidopsis* flowers does not have bracts but consist of four concentric rings of 4 sepals, 4 petals, 6 stamens and 2 fused carpels. In contrast, the primary flower of the *fif* mutant had bracts as well as sepals but the petals were incompletely developed or entirely missing (Fig. 1b, e). In addition, there were either no stamens or the stamens displaying an aberrant development (Fig. 1b, e). Furthermore, there were more than 2 carpels per flower, which were not or only partially overgrown and did not establish fertile ovaries. Most obvious, however, was the outgrowth of stems from the axillary meristems of the bracts, which carried terminal secondary flowers. A few secondary *fif* flowers showed a wild-type-like phenotype and were, thus, fertile (Fig. 1b, e).

Furthermore, the *fif* mutant plant displayed a bushy habitus compared to wild-type No-0 (Fig. 2a, b). This bushy appearance was due to an enhanced number of stem-born side branches compared to wild-type No-0, whereas the number of rosette-born side shoots was the same in *fif* and wild-type No-0 plants (Fig. 2c). In addition, *fif* mutant plant exhibited delayed flowering compared to wild-type No-0 (Fig. 2a, b).

**Figure 2.**
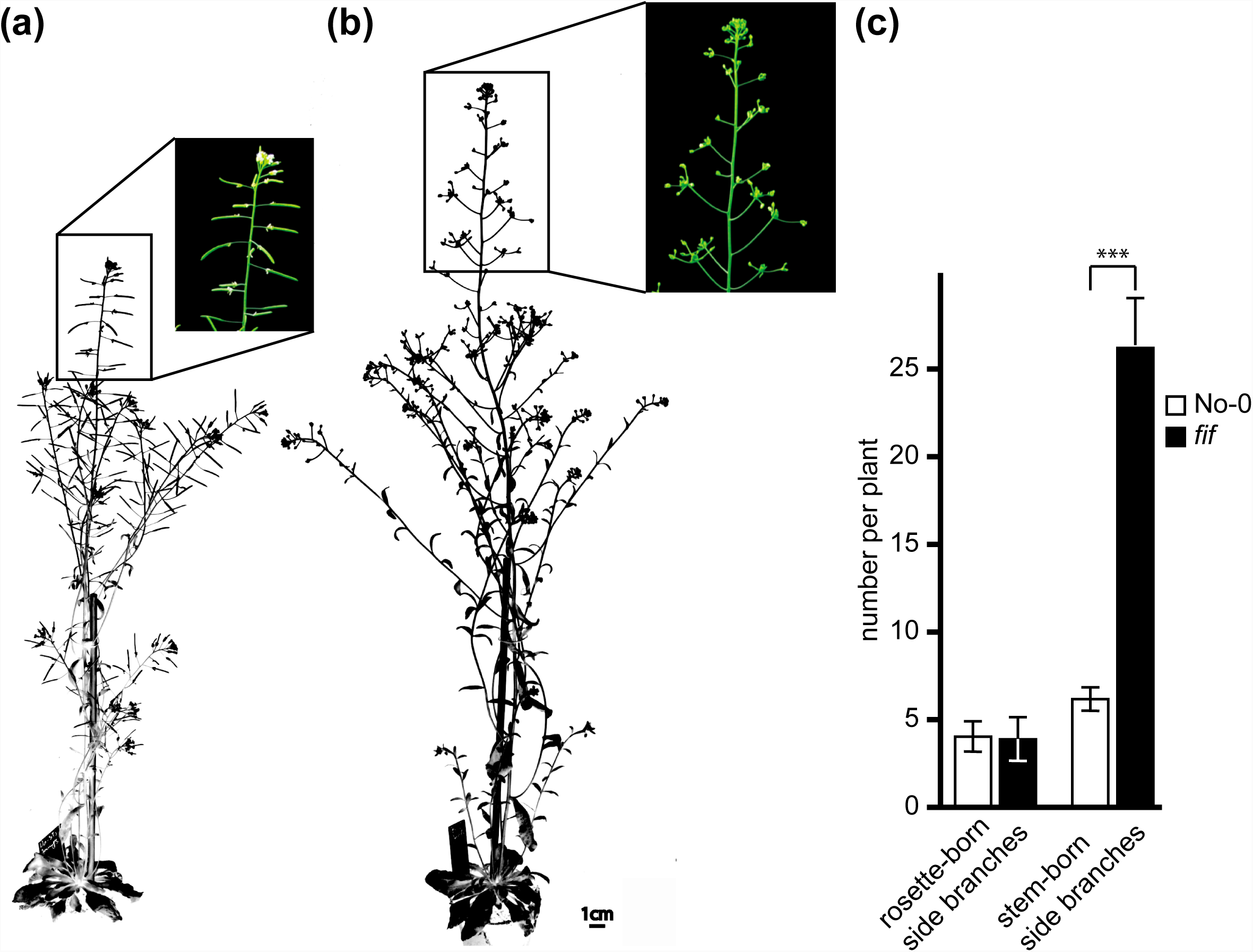
Growth habitus and degree of branching of wild-type No-0 and *fif* mutant plants. (a-b) Overview over the growth habitus and magnification of the inflorescence of 6.5-weeks old wild-type No-0 (a) and *fif* (b) plants, grown side-by-side in the greenhouse. Size bar: 1.0 cm. (c) Number of rosette-born side branches and stem-born side branches of wild-type No-0 (white bars) and *fif* (black bars) plants. Error bars indicate the standard deviation of the mean (n_No-0_ = 33, n_*fif*_ = 25, ***: p = 2x10^-23^).

### The transposon insertion is not responsible for the *fif* phenotype

According to the RIKEN RAPID and our own genotyping results, the *Ds* transposon was located in the second exon of the gene *At1g20990* that codes for a putative cysteine/histidine-rich C1 domain protein with an as yet unknown function. To validate the causal relationship between the *fif* phenotype and the *Ds* transposon insertion, we analysed an independent insertion mutant in the *Arabidopsis thaliana* Col-0 background, which exhibited a T-DNA insertion in the promoter region of *At1g20990* (SALK_073291; Alonso *et al.*, 2003). However, homozygous mutant plants of this line showed no aberrant phenotype compared to wild-type (Col-0) with respect to floral development, flower morphology, flowering time and growth habitus.

This observation raised doubts as to whether there is a functional link between the *Ds* transposon insertion and the *fif* mutant phenotype. We therefore performed a (co-) segregation analysis by backcrossing the *fif* mutant with wild-type No-0 in both directions (♀*fif* x ♂No-0, ♀No-0 x ♂*fif*). Irrespective of the direction, the crosses were successful as demonstrated by PCR on genomic DNA extracted from F1 plants using *Ds* transposon-and *At1g20990*-specific primers (Figure S1). All tested F1 plants were heterozygous for the *Ds* transposon and wild-type *At1g20990* and displayed wild-type floral organs and growth habiti (Figure S1). Therefore, the mutation that causes the *fif* phenotype is recessive. Next, six F1 plants were self-fertilized and 20 to 30 progenies each analysed for their pheno-and genotypes. As shown in figure 3, around one quarter of the F2 plants displayed the *fif* phenotype indicating that it is caused by a single mutant gene. Intriguingly, our genotyping results showed that the *Ds* transposon insertion did not co-segregate with the *fif* phenotype: 29 % of the *fif* phenotype-displaying plants did not contain the transposon, an additional 49% contained the transposon insertion only heterozygously (Figure 3). These results prove that the *Ds* insertion into the *At1g20990* locus does not cause the *fif* phenotype.

**Figure 3.**
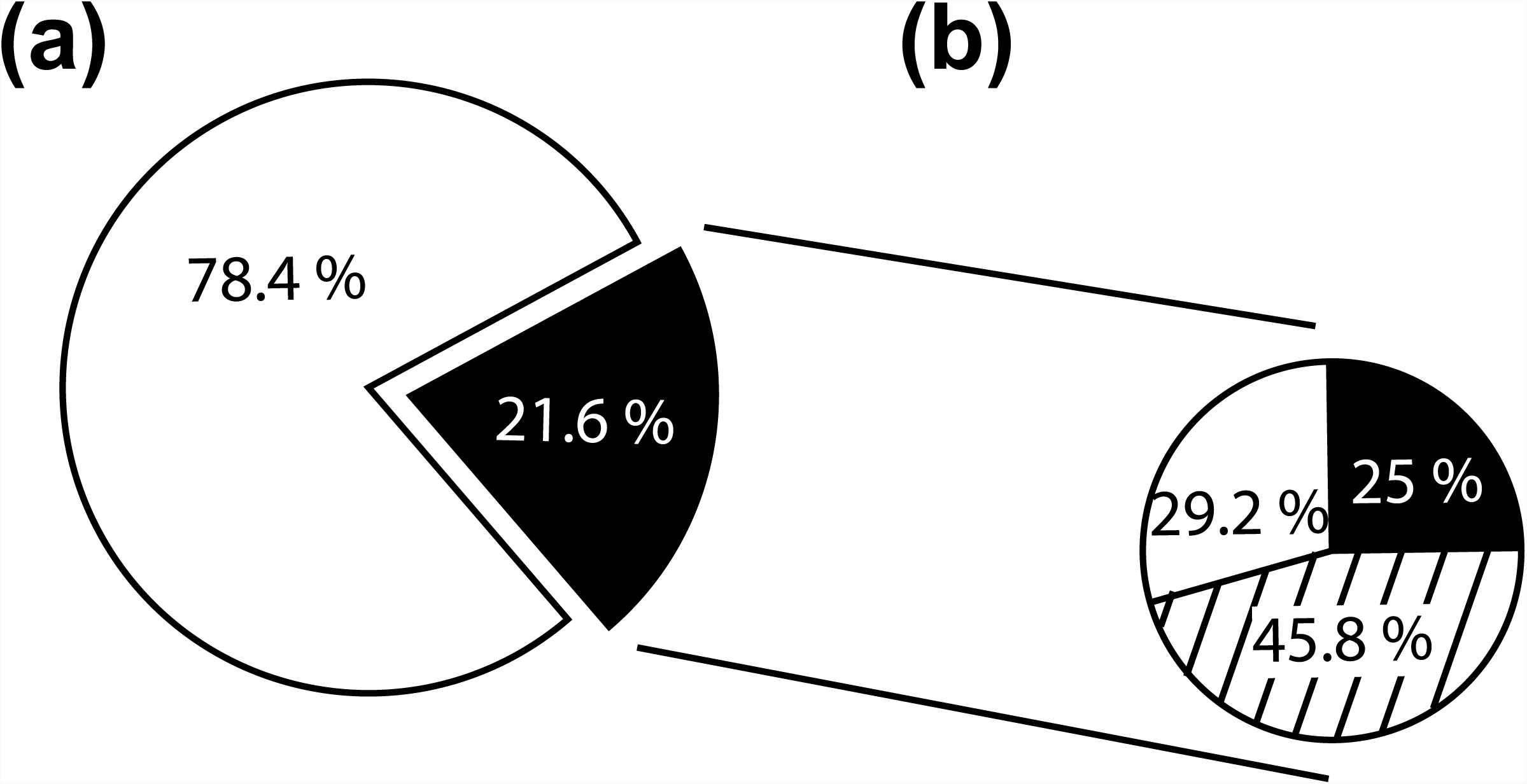
Segregation of the floral phenotype and the Ds transposon insertion within the combined F_2_ population of (♀*fif* x ♂No-0) and (♀No-0 x ♂*fif*) backcrosses. (a) Distribution of F_2_ plants, showing either the wild-type (78.4 %) or the *fif* floral phenotype (21.6 %). (b) Distribution of the transposon insertions within the plants of the F_2_ population that displayed the *fif* floral phenotype. White circle outcut: no transposon insertion (29.2 %), striped outcut: heterozygous for the Ds transposon insertion (45.8 %), black outcut: homozygous for the Ds transposon insertion (25.0 %).

### The *fif* phenotype is caused by a novel allele of *LEAFY* (*LFY*)

To identify the mutant locus genetically responsible for the *fif* phenotype, we combined a classical mapping (Neff *et al.*, 2002; Kover *et al.*, 2009; Pacurar *et al.*, 2012) with a mapping-by-sequencing approach (James *et al.*, 2013; Schneeberger, 2014). To establish a mapping population, *fif* mutant plants (No-0) were crossed in both direction with plants of the Col-0 accession. Irrespective of the crossing direction, all the F1 plant displayed a wild-type phenotype (Figure S2a). Eight F1 plants were self-fertilized and 1582 F2 plants characterized phenotypically. In accordance with the self-crossing results described above, around 25 % of the F2 plants (437 of the 1582) showed the *fif* phenotype (Figure S2b). Leaf material was harvested from 425 of the 437 F2 plants in groups of 15 to 20 individuals; in addition leaf material from 200 F2 plants was collected individually. Genomic DNA was extracted and used for classical mapping. Using chromosome-specific INsertion and DELetion (INDEL) markers (Pacurar *et al.*, 2012) the mutant locus was mapped to the q-arm of chromosome 5 (Figure 4a). Two additional INDEL markers and two Single Nucleotide Polymorphism (SNP) based Derived Cleaved Amplified Polymorphic Sequences (dCAP) markers (Kover *et al.*, 2009; Neff *et al.*, 2002) limited the Quantitative Trait Locus (QTL) responsible for the *fif* phenotype to the terminal end of chromosome 5’s q-arm (Figure 4b, dCAP S5-24: 99% No-0).

**Figure 4.**
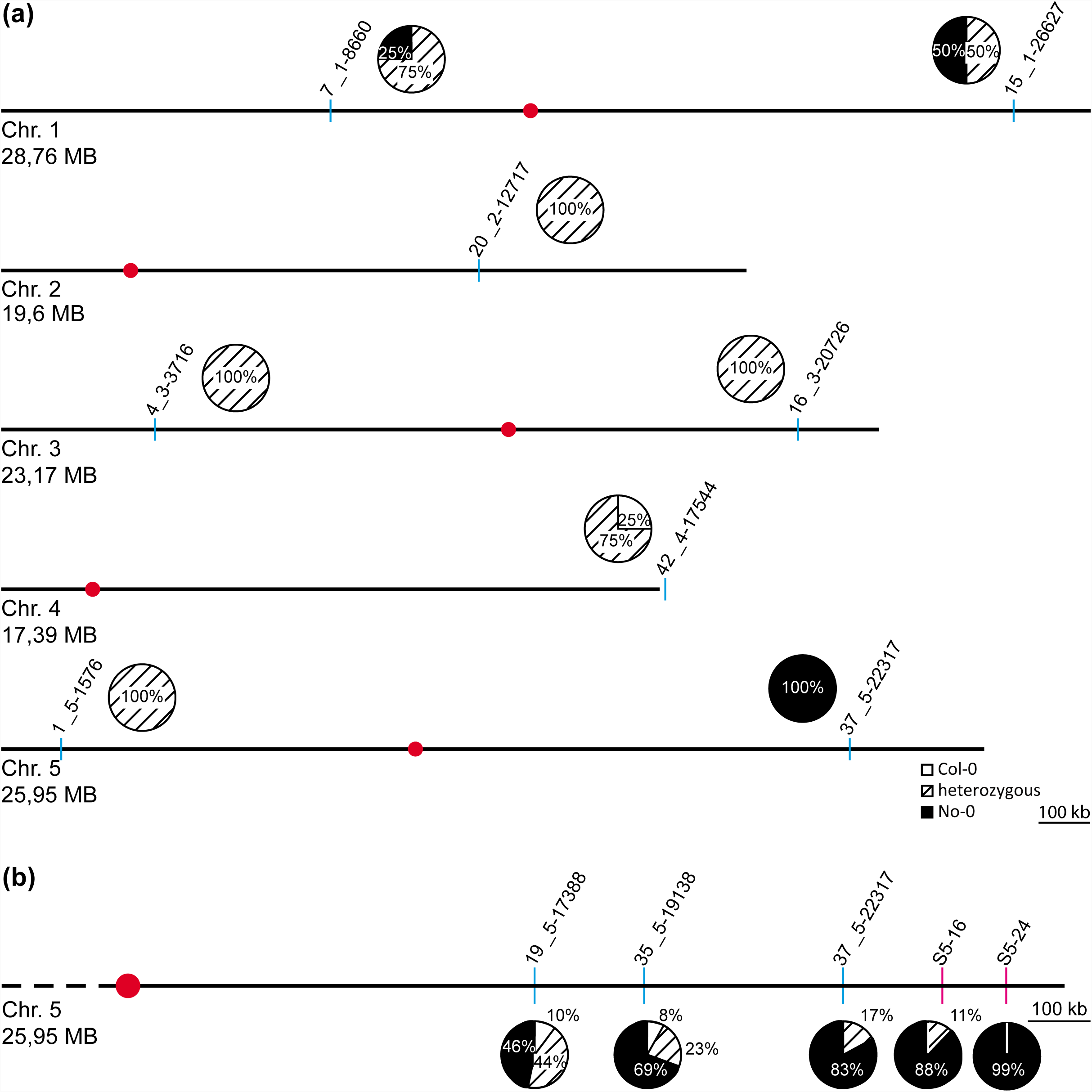
INDEL marker-and SNP-based dCAP marker-associated containment of the *fif* locus using a mapping population generated by a cross of the *fif* mutant (No-0) with wild-type Col-0. (a) Schematic representation of the 5 *A. thaliana* chromosomes (sizes in MB) and the localization of the chromosome-specific INDEL markers initially used for mapping (codes above blue lines). (b) Schematic representation of the q-arm of chromosome 5 and the localization of INDEL (codes above the blue lines) and SNP-based dCAP markers (codes above red lines) used for fine mapping. The pie charts show the distribution of the No-0 and Col-0 genotypes for each chromosome (a) and the q-arm of chromosome 5 (b). White circular outcut: homozygous for Col-0, striped outcut: heterozygous for Col/No-0, black outcut: homozygous for No-0; red dot: localization of the centromere.

To establish the exact localization of the mutant locus, we deep-sequenced the total genome of 245 homozygous *fif* mutant plants derived from the *fif* (No-0) x WT (Col-0) crosses described above, and determined the frequencies of No-0 and Col-0 alleles along the chromosomes. Whereas the heterozygous distribution of No-0 and Col-0 sequences was found to be equal with respect to chromosomes 1 to 4 (Figure S3a-d), there was a very significant deviation towards No-0 sequences at the terminal end of chromosome 5 (Figure 5a). A detailed examination of this 300 kb stretch revealed 100 % identity with the No-0 sequence (Figure 5b). This sequence stretch conformed with the QTL identified by classical mapping.

**Figure 5.**
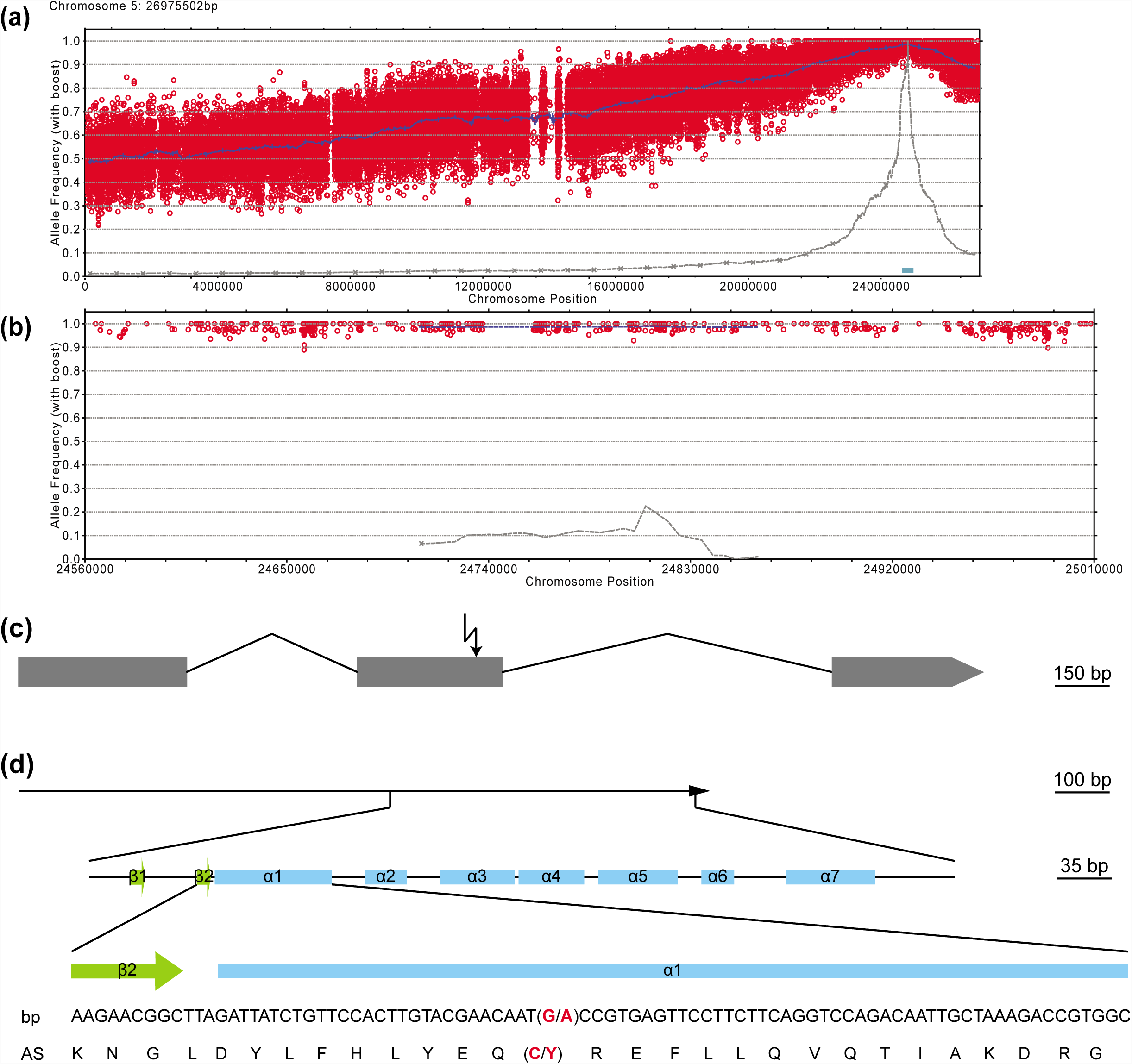
Identification of the *fif*-related SNP in the second exon of the *LEAFY* (*LFY*) locus on chromosome 5 by genome sequencing of a mapping population generated by a cross of the *fif* mutant (No-0) with wild-type Col-0. (a) Allele frequency analysis of the Nos genotype within chromosome 5 of the recombinant mutant pool. Each red circle refers to a SNP marker distinguishing the Nos and Col genotypes. The blue line refers to a 200 kb sliding window analysis of the allele frequencies. The brown line and blue box highlight the estimated mapping intervals (x-axis: genomic location; y-axis: Nos allele frequency). (b) Like (a), but only showing the 300 kb mapping interval. (c) Exon-intron organization of the *LFY* locus with the *fif*-related SNP marked by an arrow. Exons are shown as grey boxes and introns as exons connecting lines. (d) Sequence of the *LFY* gene showing the *fif* SNP (G to A exchange, red) and the resulting amino acid exchange (C to Y, red) within the DNA-binding domain of the LFY protein. Green boxes: β-sheets; blue boxes: α-helices (according to Hames et al., 2008).

A detailed comparison of the *fif* and wild-type No-0 sequence in this 300 kb stretch revealed a single SNP, which did not result in a silent mutation but caused a change in a codon. This SNP was also found in all the 143 individually tested *fif* mutant plants and reflected a single guanine-to-adenine exchange in the second exon of the *LEAFY* (*LFY*) gene (*At5g61850*, Figure 5c). This mutation caused a cysteine-to-tyrosine amino acid exchange at position 263 in the DNA-binding domain of the LFY protein (Figure 5d). To prove that this point mutation causes the *fif* phenotype, we transformed the *fif* mutant (No-0) with constructs expressing LFY-GFP or LFY^FIF^-GFP under the control of the *35S* promoter. Whereas the expression of LFY-GFP complemented the *fif* mutant phenotype almost completely, there was no complementation with LFY^FIF^-GFP (Figure S4).

### LFY^FIF^ impairs DNA-binding capability but shows wild-type intracellular localization and homomerization

Having identified a new *LFY* allele to be responsible for the *fif* phenotype, we next analysed the putative consequences of the Cys263-to-Tyr exchange for LFY protein properties at molecular and cell biological levels.

To test a putative alteration in subcellular localization, C-terminal GFP fusions of wild-type LFY and the mutant LFY version (LFY^FIF^) were expressed under the control of the *Arabidopsis ubiquitin 10* (*UBQ10*) promoter in transiently transformed *Nicotiana benthamiana* epidermal leaf cells. The functionality of C-(and N-terminal) GFP fusions of LFY was previously shown by the genetic complementation of the *lfy-12* mutant phenotype (Wu *et al.*, 2003). As shown in figure 6a, LFY-GFP and LFY^FIF^-GFP localised to the cytoplasm and the nucleus in a similar manner. The observed fluorescence pattern of LFY-GFP and LFY^FIF^-GFP is in accordance with the pattern previously reported for their expression in tobacco epidermal leaf cells (Siriwardana and Lamb, 2012b).

**Figure 6.**
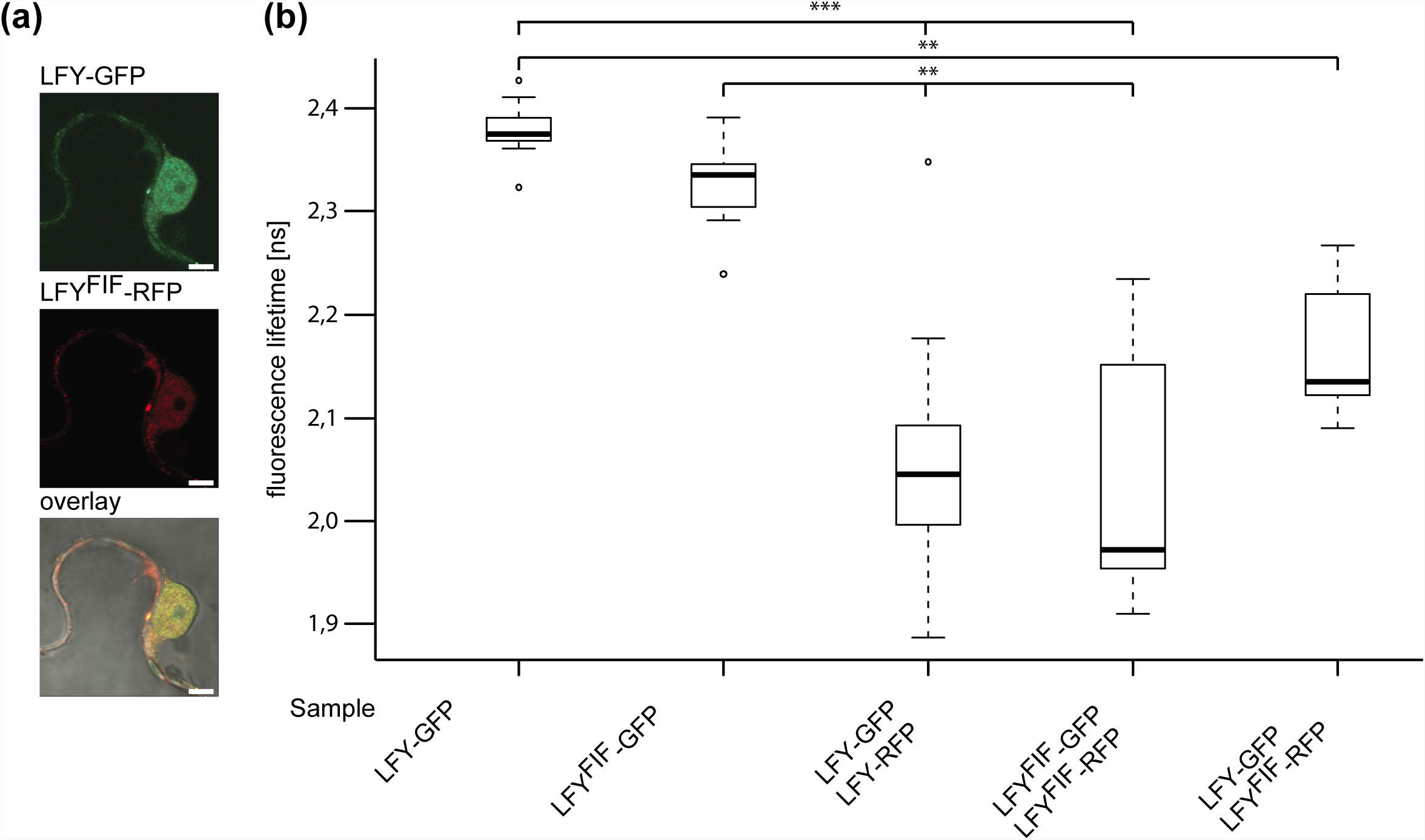
Comparative analysis of the intracellular localization and homomerization capacity of LFY and LFY^FIF^. (a) Confocal fluorescence images of transiently transformed *Nicotiana benthamina* epidermal leaf cells expressing LFY-GFP and LFY^FIF^-RFP in the same cell. Size bar: 5 µm. (b) FRET-FLIM analysis of the homo-and heterotypic interaction of LFY and LFY^FIF^. LFY-GFP or LFY^FIF^-GFP were expressed either alone or together with the indicated RFP fusions and the fluorescence lifetime of the GFP fusions measured in nucleus. A reduction of the GFP fluorescence lifetime indicates interaction. The data are presented in Box-and-Whisker plots including the median (thick line), the upper and lower quartile (+/-25%, white boxes), the maximum and minimum (dottet line) and outlier points (n > 20, each). The variance was analyzed by a Levene test and statistical significance was determined with an all-pair, two-sided Kruskal-Walles test followed by an all-pair Steel-Dwass test (**: p < 0.01; ***: p < 0.001).

Next, we tested by *in vivo* FRET-FLIM whether LFY protein-protein interaction, here especially LFY homomerization (Sirwardana and Lamb, 2012a), was altered. To do so, C-terminal GFP fusions (FRET donor) and C-terminal RFP fusions (FRET acceptor) were transiently expressed, either individually (donor only) or in combination in *N. benthamiana* epidermal leaf cells and the fluorescence lifetime of the donor fusion was measured. As shown in figure 6b, the fluorescence lifetimes of LFY-GFP and LFY^FIF^-GFP were similar in the absence of the acceptor fusions. However, the lifetimes of LFY-GFP and LFY^FIF^-GFP decreased significantly when they were co-expressed with either LFY-RFP or LFY^FIF^-RFP demonstrating homotypic (LFY-LFY, LFY^FIF^-LFY^FIF^) and heterotypic (LFY-LFY^FIF^) homomerization *in planta* (Figure 6b). In addition, there was no significant difference in the interaction of the homotypic and heterotypic homomers (Figure 6b).

The Cys263-to-Tyr exchange is located in the first α-helix of the LFY DNA-binding domain (Figure 5d). We, therefore, used a quantitative DNA-protein interaction ELISA approach (qDPI-ELISA; Fischer, Böser *et al.*, 2016) to test whether the mutation interferes with the DNA-binding capability of LFY *in vitro*. We expressed N-terminally GFP-tagged full-length LFY, as well as full-length LFY^FIF^ and GFP, in *E. coli* independently and applied the crude extracts containing the fusion proteins or GFP, in identical amounts, based on the GFP fluorescence and western-blotting, to ELISA plates in two dilutions. The plates were covered with double-stranded (ds) DNA oligonucleotides representing either the LFY-binding sequence of the *AP1* promoter (*pAP1*), a mutated *pAP1* version (*pAP1m*) that is not recognized by LFY (Winter *et al.*, 2011) a random sequence without any similarity to the LFY binding motif (*C28M12*), or were uncovered. The DNA-binding efficiency of the proteins was recorded by determining the GFP fluorescence of the bound proteins (Fischer, Böser *et al.*, 2016). GFP-LFY exhibited a specific binding to *pAP1* and no binding to any other oligonucleotide or to the oligonucleotide-free ELISA plate (Figure 7). In contrast, GFP-LFY^FIF^, like GFP or the *E. coli* crude extract without recombinant protein, was unable to recognize *pAP1* or any other oligonucleotide (Figure 7). To exclude the possibility that the Cys263-to-Tyr exchange may alter the DNA-binding specificity we used a DPI-ELISA based approach to screen a dsDNA oligonucleotide library reflecting 4096 randomized DNA hexamers (Brand *et al.*, 2013a, b) with GFP-LFY-and GFP-LFY^FIF^-containing *E.coli* extracts. Whereas a DNA-binding consensus sequence was obtained for GFP-LFY (5’-GGGC-3’/3’-CCCG-5’), there was no DNA-binding of GFP-LFY^FIF^ to any oligonucleotide in the library.

**Figure 7.**
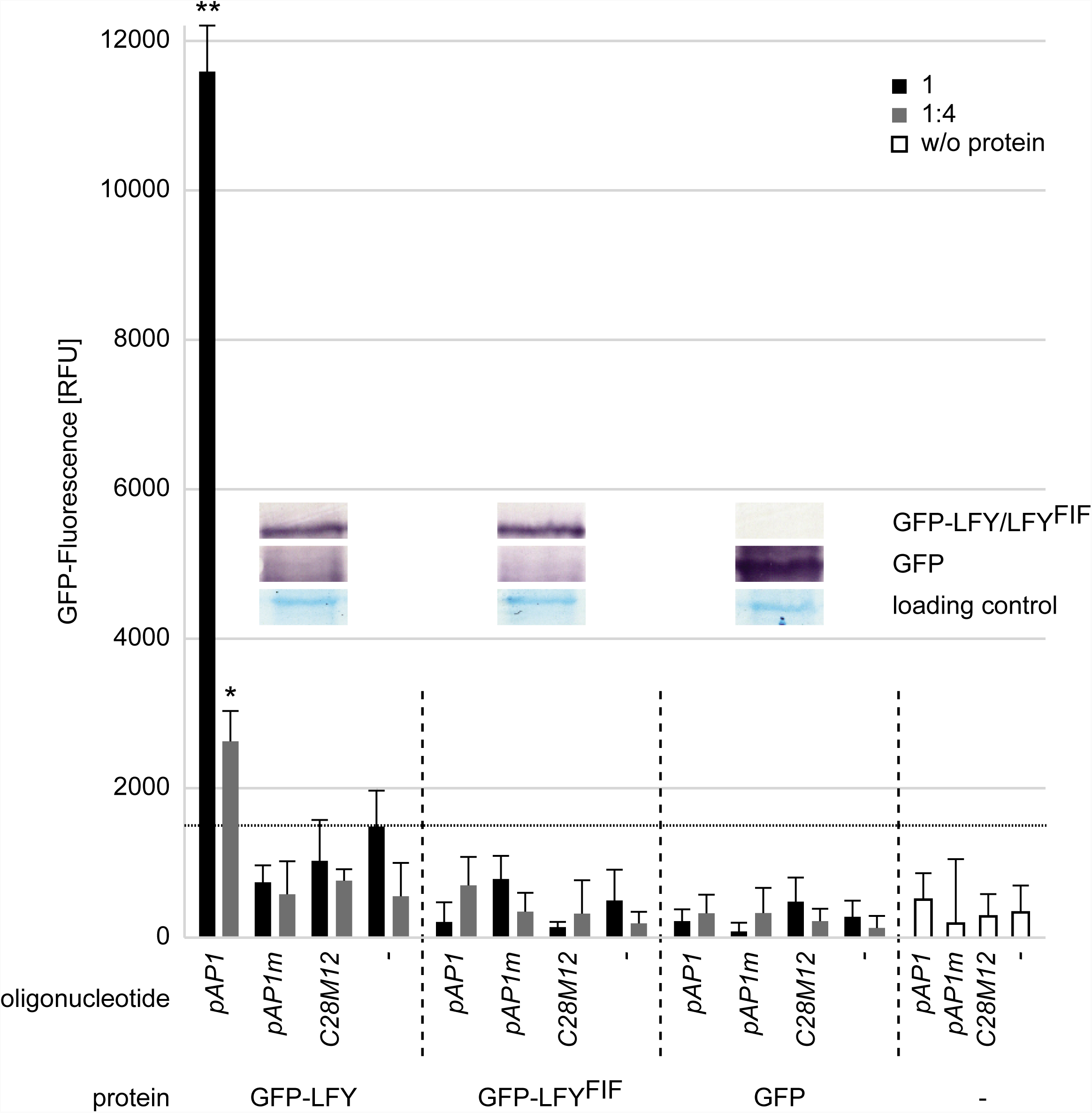
Comparative analysis of the *in vitro* DNA-binding capacity of LFY and LFY^FIF^ using a GFP-fluorescence-based DPI-ELISA approach. GFP-LFY and GFP-LFYFIF were expressed in *E.coli*. After extraction, crude extracts containing either no recombinant protein (w/o protein) or, based on GFP fluorescence, equal amounts of GFP or GFP fusion protein were added to ELISA plates covered with either the double-stranded (ds) DNA oligonucleotide *pAP1*, which contains a LFY recognition site, an altered version of *pAP1* (*pAP1m*), in which the recognition site was mutated, a dsDNA oligonucleotide unrelated to the *pAP1* and *pAP1m* sequences (*C28M12*) or without any DNA-oligonucleotide. The amount of DNA-bound fusion protein was detected by reading out the GFP fluorescence. The crude extract was either used undiluted (black bars) or in a 1:4 dilution (grey bars). Error bars indicate the standard deviation of the mean (n = 3) and asteriks statistically significant differences to the background fluorescence (dotted horizontal line), determined by two-sided t-test (*: p < 0.05; **: p < 0.01). The inlet shows a Western-blot of the crude extracts using a GFP polyclonal antiserum for detection of GFP, GFP-LFY and GFP-LFY^FIF^ as well as a Coomassie stain as loading control.

## Discussion

In our search for novel floral genes in *Arabidopsis thaliana* we identified the *fif* Ds transposon insertion mutant in the No-0 accession in the RIKEN RAPID collection (Ito *et al.*, 2002; Kuromori *et al.*, 2004). *fif* mutant plants display a novel floral phenotype and inflorescence architecture, as they develop aberrant and infertile primary flowers in combination with short stems that emerge from vegetative meristems in the axillars of the bracts and carry fertile secondary flowers.

The *Ds* transposon insertion in the genome of the *fif* mutant was annotated to gene *At1g20990*, which encodes a cysteine/histidine-rich C1 domain protein. However, as demonstrated by our genetic analysis, the *Ds* transposon insertion into the *At1g20990* locus is not the cause of the *fif* phenotype. Obviously, another mutant locus generated somewhere else in the genome, most likely during transposon movement, is responsible for the *fif* phenotype. Using combined classical and genome sequencing-based mapping approaches, the causal mutation for the *fif* phenotype was found to be in the *LFY* gene. The mutation is a single G-to-A exchange in the second exon of *LFY*, creating the novel, recessive *lfy* allele. The mutation causes a Cys-to-Tyr exchange at position 263 in the LFY^FIF^ amino acid sequence.

The cell biological analysis of LFY-GFP and LFY^FIF^-GFP revealed an intracellular localization in the cytoplasm and nucleus of tobacco epidermal leaf cells identical to that previously reported for LFY-GFP (Siriwardana and Lamb, 2012a). Thus, a mis-localisation cannot be the cause of the LFY^FIF^ malfunction. In addition, as shown by quantitative FRET-FLIM interaction studies the mutation does not interfere with the homomerization capacity of LFY. Especially the latter result was to be expected as the domain essential for homomerization is located at the N-terminus of LFY (amino acid 46 to 127; Siriwardana and Lamb, 2012a).

However, our quantitative DPI-ELISA assay demonstrated that, in contrast to LFY-GFP, LFY^FIF^-GFP lost its capacity to bind to its DNA target, as it is present, for instance, in the *AP1* promoter (Winter *et al.*, 2011). Furthermore, the DPI-ELISA based approach for the determination of putative alterations in binding specificity did not reveal any DNA-binding activity for LFY^FIF^-GFP.

According to the available crystal structure of the DNA-bound dimer, Cys263 is well conserved between the LFY homologs of many plant species but has never previously been reported to be crucial for DNA-binding (Hames *et al.*, 2008). Intriguingly, Cys263 does not contribute to the physical contact of LFY with DNA; however, the α1-helix, in which Cys263 is positioned, participates in the cooperative DNA-binding of LFY, as it facilitates the establishment and stabilization of the DNA-binding domains in the minor and major grove of DNA (Hames *et al.*, 2008). Therefore, the change of the relatively small Cys to the bulky, aromatic Tyr might prevent the folding of the α1-helix and thereby strongly restrict the cooperative binding of LFY to its target DNA.

The total failure of LFY^FIF^ to bind to DNA explains the strong floral phenotype of especially the primary flowers. LFY is one of the master regulators in the FLIP of *Arabidopsis* (and other plant species) and controls, together with other factors and *via* a complex regulatory network, the spatiotemporal expression of downstream FLIP genes and also of the homeotic flower genes required for flower organ formation. Although only a single amino acid exchange is affected, LFY^FIF^ mirrors in principle the flower phenotype of known strong *lfy* alleles. However, of the more than 15 described *lfy* alleles (Weigel *et al.*, 1992), the six alleles that show such a strong floral phenotype produce shortened LFY polypeptides caused by either premature stop codons (*lfy-1, lfy-6, lfy-7, lfy-8, lfy-11*) or a non-sense frame shift C-terminal of Gln196 (*lfy-15*). Hence, the strong phenotype of the *fif* allele needs a different explanation: LFY^FIF^ may act dominant-negatively by either forming non-functional heteromers with wild-type LFY, which cannot longer bind to DNA, or by titrating out interaction partners required for LFY function (Siriwardana and Lamb, 2012b). However, as long as sufficient wild-type LFY is present in heterozygous plants, the *fif* mutant shows recessive inheritance.

The failure of LFY^FIF^ to bind to DNA is also explains the bushy growth architecture of the *fif* mutant. It has recently been shown (Chahtane *et al.*, 2013) that mutations in *lfy* can cause the emergence of axillary meristems instead of floral meristems resulting in an enhanced number of side branches. In addition, the ectopic expression of a nearly full-length LFY version with weaker *in vitro* DNA-binding capacity and dramatically reduced *in vivo* transcriptional activity [LFY_HARA(Δ40)_] in the Col-0 accession causes a bushy phenotype similar to that of the No-0 *fif* mutant (Chahtane *et al.*, 2013). Interestingly, the His387-to-Ala and Arg390-to-Ala in LFY_HARA(Δ40)_ are also mooted to interfere with the cooperative binding of LFY to its target DNA as well. Taken together, our data demonstrate the general importance of Cys263 for LFY function not only in floral development but also in axillary meristem outgrowth in *Arabidopsis*.

Most intriguingly, the *fif* floral phenotype appears to be specific for the No-0 accession, as, to our knowledge, it has never been reported for the Col-0 or any other accession. However, the *fif* phenotype also becomes also manifest in the Col-0 accession when the *fif* locus of No-0 is transferred to Col-0. This phenomenon might be explained by differences in the spatio-temporal transcriptional activity of the No-0 and Col-0 *LFY* loci during vegetative meristem and floral development. Therefore, the *fif* phenotype may only be visible in other accessions such as Col-0 when the No-0 locus is artificially introduced into them and drives LFY^FIF^ accumulation.

## Experimental procedures

### Plant material

Seeds of the homozygous *Ds* transposon insertion line 15-3794-1 and the corresponding wild-type accession (No-0) were obtained from the RIKEN *Arabidopsis* Phenome Information database (RAPID; Kuromori *et al.*, 2006). Seeds of the homozygous T-DNA insertion line Salk_073291 and the corresponding wild-type accession (Col-0) were obtained from the Nottingham *Arabidopsis* Stock Centre (NASC; Alonso *et al.*, 2003).

### Plasmid construction

Using gene-specific primers [sense (S): 5’-caccATGGATCCTGAAGGTTTCACG-3’, antisense (A): 5’-GAAACGCAAGTCGTCGCCG-3’) the cDNA of *LFY* was amplified from pSST14 (gift Jan Lohmann, University of Heidelberg, Germany) and cloned in pENTR™ /D-TOPO^®^. Site-directed mutagenesis (SDM) was performed to produce the *fif* cDNA using the following primers (S: 5’-CTGTTCCACTTGTACGAACAATaCCGTGAGTTCCTTCTTCAG-3’, A: 5’-CTGAAGAAGGAACTCACGGtATTGTTCGTACAAGTGGAACAG-3’). With Gateway™ LR Clonase™ II Enzyme mix the *LFY* cDNA was inserted into pUGT1-Dest (A. Hahn, unpublished) and pB7RWG2-Dest (Karimi *et al.*, 2002) for plant expression and into pET-Dest42GFP (Fischer, Böser *et al.*, 2016) for *E. coli* expression.

### Classical mapping and mapping by genome sequencing

Genetic mapping was accomplished using 100 phenotypic *fif* plants collected from a F2 population derived from a cross between *fif* (No-0) and Col-0. The mapping strategy and the molecular markers used to identify the causal locus were described by Păcurar *et al.* (2012). After mapping of the chromosome arm and next-generation sequencing (NGS, see below) the point mutation was confirmed by derived cleaved-amplified polymorphic sequence primers designed by using the dCAPS Finder 2.0 software (Neff, Turk and Kalishman, 2002). One or two mismatches were introduced in one of the used primer to incorporate an allele-specific restriction site into the PCR product. After amplification, the PCR products were digested (enzymes from Thermo Scientific) following the manufacturer’s recommendations and separated on a 4% agarose gel. All used markers are listed in table S1.

NGS mapping was performed using a pool of 425 phenotypic *fif* plants from the crossing described above. A pool of 40 wild-type No-0 plants was sequenced to generate a genome-wide marker list and to mine the *fif* genome for acquired mutations. Isolation of genomic DNA was performed in groups up to 20 plants using the DNeasy^®^ Plant Mini Kit (QIAGEN) following the manufacturer’s recommendations. DNA concentration was determined with the use of NanoDrop ND-1000 and the whole pool composed by using 100 µg DNA of each group. Sequencing was performed at the Max Planck-Genome-Centre Cologne by a HiSeq2500 (Illumina) Sequencer producing ∼35.000.000 read-pairs for each pool. Short reads of both pools were respectively aligned against the Col-0 reference sequence (TAIR10) and SNPs were called using *shore* pipeline (version v0.8) with *GenomeMapper* (version v0.4.4s) with default parameters (Ossowski *et al.*, 2008; Schneeberger *et al.*, 2009a). Genome-wide SNP markers were defined with filtering for sequencing coverage and allele frequency using *SHOREmap* (version 3.0, Sun *et al.*, 2015; Schneeberger *et al.*, 2009b; Schneeberger, 2014). Sliding window-based estimation of allele frequencies of the Nos allele in the pooled F2 samples and identification of a mapping interval were performed with *SHOREmap* (version 3.0) using default parameters. Comparison of the consensus calls of both pools in the 300 kb mapping interval revealed the mutation in *LFY*.

### Localization and FRET-FLIM studies

The indicated constructs and p19 as gene silencing suppressor were transformed into *Agrobacterium tumefaciens* strain GV3101 and infiltrated into *Nicotiana benthamiana* leaves. The localization of the fusion proteins was performed 3 days after infiltration using 488 nm or 561 nm lasers for GFP or RFP excitation, respectively, at the SP8 laser scanning microscope (Leica Microsystems GMBH) with LAS AF and SymPhoTime software using a 63x/1.20 water immersion objective (Ladwig *et al.*, 2015). FLIM data were derived from measurements of at least 20 probes for each fusion protein combination. To excite LFY-GFP and LFY^FIF^-GFP for FLIM experiments, a 470 nm pulsed laser (LDH-P-C-470) was used, and the corresponding emission was detected with a SMD Emission SPFLIM PMT from 495 to 545 nm by time-correlated single-photon counting using a Picoharp 300 module (PicoQuant). Each time-correlated single-photon counting histogram was reconvoluted with the corresponding instrument response function and fitted against a monoexponential decay function for donor-only samples and a biexponential decay function for the other samples to unravel the GFP fluorescence lifetime of each probe. The average GFP fluorescence lifetimes as well as the standard error values were calculated using Microsoft Excel 2013. To test for homogenity of variance Levene’s test (df=5/140, F=26.298, p < 0.0001) was used and statistical significance was calculated by a two-tailed, all-pair Kruskal-Wallis test followed by a Steel-Dwass *post hoc* correction using JMP version 12.2.0 (Ohmi *et al.*, 2016).

### qDPI-ELISA, DPI-ELISA based screening and western blotting

qDPI-ELISA was performed using *E.coli* crude extracts containing GFP-tagged LFY or LFY^FIF^, GFP alone or no fluorescent protein according to Fischer, Böser *et al.* (2016). The sequences of the 5’-biotinylated dsDNA oligonucleotides *AP1, mAP1* and *C28M12* used for the immobilization on Streptavidin-coated 384 well microtiter plate are displayed in table S2. Before addition to the microtiter plate, the equal content of GFP-tagged fusion protein in the crude extracts was adjusted according to the GFP fluorescence using a fluorescence reader (TECAN Safire).

The DPI-ELISA based specificity screening, using a dsDNA oligo array on a 384 well microtiter plate covering all possible 4096 hexanucleotide DNA motifs was performed as described previously (Brand *et al.* 2013a, b).

## Acknowledgements

The authors would like to thank J. Lohmann (Universität Heidelberg, Germany) for the *LFY* cDNA, M. Fischer for technical support, J. Schröter and G. Huber for support in plant cultivation, F. de Courcy for proofreading the manuscript and the members of the multidisciplinary graduate school “Morphological Variability of Organisms under Environmental Stress” for discussion. This work was supported by a grant of the Ministerium für Wissenschaft, Forschung und Kunst Baden-Württemberg to A. Mohrholz.

## Supporting information

Additional Supporting Information is found in the on-line version of this article.

## References

Alonso, J.M., Stepanova, A.N., Leisse, T.J., Kim, C.J., Chen, H., Shinn, P., Stevenson, D.K., Zimmerman, J., Barajas, P., Cheuk, R., Gadrinab, C., Heller, C., Jeske, A., Koesema, E., Meyers, C.C., Parker, H., Prednis, L., Ansari, Y., Choy, N., Deen, H., Geralt, M., Hazari, N., Hom, E., Karnes, M., Mulholland, C., Ndubaku, R., Schmidt, I., Guzman, P., Aguilar-Henonin, L., Schmid, M., Weigel, D., Carter, D.E., Marchand, T., Risseeuw, E., Brogden, D., Zeko, A., Crosby, W.L., Berry, C.C. and Ecker, J.R. (2003) Genome-wide insertional mutagenesis of Arabidopsis thaliana. Science, 301, 653–657.

Blazquez, M.A., Ferrandiz, C., Madueno, F. and Parcy, F. (2006) How floral meristems are built. Plant Mol Biol, 60, 855–870.

Bowman, J.L., Alvarez, J., Weigel, D., Meyerowitz, E.M. and Smyth, D.R. (1993) Control of flower development in Arabidopsis thaliana by APETALA1 and interacting genes. Development, 119, 721–743.

Brand, L.H., Henneges, C., Schussler, A., Kolukisaoglu, H.U., Koch, G., Wallmeroth, N., Hecker, A., Thurow, K., Zell, A., Harter, K. and Wanke, D. (2013a) Screening for protein-DNA interactions by automatable DNA-protein interaction ELISA. PLoS One, 8, e75177.

Brand, L.H., Fischer, N.M., Harter, K., Kohlbacher, O. and Wanke, D. (2013b) Elucidating the evolutionary conserved DNA-binding specificities of WRKY transcription factors by molecular dynamics and in vitro binding assays. Nucleic Acids Res, 41, 9764–9778.

Chahtane, H., Vachon, G., Le Masson, M., Thevenon, E., Perigon, S., Mihajlovic, N., Kalinina, A., Michard, R., Moyroud, E., Monniaux, M., Sayou, C., Grbic, V., Parcy, F. and Tichtinsky, G. (2013) A variant of LEAFY reveals its capacity to stimulate meristem development by inducing RAX1. Plant J, 74, 678–689.

Coen, E.S. and Meyerowitz, E.M. (1991) The war of the whorls: genetic interactions controlling flower development. Nature, 353, 31–37.

Fischer, S.M., Böser, A., Hirsch, J.P. and Wanke, D. (2016) Quantitative Analysis of Protein-DNA Interaction by qDPI-ELISA. Methods Mol Biol, 1482, 49–66.

Hames, C., Ptchelkine, D., Grimm, C., Thevenon, E., Moyroud, E., Gerard, F., Martiel, J.L., Benlloch, R., Parcy, F. and Muller, C.W. (2008) Structural basis for LEAFY floral switch function and similarity with helix-turn-helix proteins. EMBO J, 27, 2628–2637.

Huala, E. and Sussex, I.M. (1992) Leafy Interacts with Floral Homeotic Genes to Regulate Arabidopsis Floral Development. Plant Cell, 4, 901–913.

Irish, V.F. and Sussex, I.M. (1990) Function of the Apetala-1 Gene during Arabidopsis Floral Development. Plant Cell, 2, 741–753.

Ito, T., Motohashi, R., Kuromori, T., Mizukado, S., Sakurai, T., Kanahara, H., Seki, M. and Shinozaki, K. (2002) A new resource of locally transposed Dissociation elements for screening gene-knockout lines in silico on the Arabidopsis genome. Plant Physiol, 129, 1695–1699.

James, G.V., Patel, V., Nordstrom, K.J.V., Klasen, J.R., Salome, P.A., Weigel, D. and Schneeberger, K. (2013) User guide for mapping-by-sequencing in Arabidopsis. Genome Biol, 14.

Karimi, M., Inze, D. and Depicker, A. (2002) GATEWAY vectors for Agrobacterium-mediated plant transformation. Trends Plant Sci, 7, 193–195.

Kempin, S.A., Savidge, B. and Yanofsky, M.F. (1995) Molecular-Basis of the Cauliflower Phenotype in Arabidopsis. Science, 267, 522–525.

Kover, P.X., Valdar, W., Trakalo, J., Scarcelli, N., Ehrenreich, I.M., Purugganan, M.D., Durrant, C. and Mott, R. (2009) A Multiparent Advanced Generation Inter-Cross to fine-map quantitative traits in Arabidopsis thaliana. PLoS genetics, 5, e1000551.

Kuromori, T., Hirayama, T., Kiyosue, Y., Takabe, H., Mizukado, S., Sakurai, T., Akiyama, K., Kamiya, A., Ito, T. and Shinozaki, K. (2004) A collection of 11 800 single-copy Ds transposon insertion lines in Arabidopsis. Plant J, 37, 897–905.

Kuromori, T., Wada, T., Kamiya, A., Yuguchi, M., Yokouchi, T., Imura, Y., Takabe, H., Sakurai, T., Akiyama, K., Hirayama, T., Okada, K. and Shinozaki, K. (2006) A trial of phenome analysis using 4000 Ds-insertional mutants in gene-coding regions of Arabidopsis. Plant J, 47, 640–651.

Ladwig, F., Dahlke, R.I., Stuhrwohldt, N., Hartmann, J., Harter, K. and Sauter, M. (2015) Phytosulfokine Regulates Growth in Arabidopsis through a Response Module at the Plasma Membrane That Includes CYCLIC NUCLEOTIDE-GATED CHANNEL17, H+-ATPase, and BAK1. Plant Cell, 27, 1718–1729.

Lohmann, J.U. and Weigel, D. (2002) Building beauty: the genetic control of floral patterning. Dev Cell, 2, 135–142.

Ma, H. (1998) To be, or not to be, a flower--control of floral meristem identity. Trends Genet, 14, 26–32.

Mandel, M.A., Gustafsonbrown, C., Savidge, B. and Yanofsky, M.F. (1992) Molecular Characterization of the Arabidopsis Floral Homeotic Gene Apetala1. Nature, 360, 273–277.

Miwa, H., Betsuyaku, S., Iwamoto, K., Kinoshita, A., Fukuda, H. and Sawa, S. (2008) The Receptor-Like Kinase SOL2 Mediates CLE Signaling in Arabidopsis. Plant Cell Physiol, 49, 1752–1757.

Neff, M.M., Turk, E. and Kalishman, M. (2002) Web-based primer design for single nucleotide polymorphism analysis. Trends Genet, 18, 613–615.

Ohmi, Y., Ise, W., Harazono, A., Takakura, D., Fukuyama, H., Baba, Y., Narazaki, M., Shoda, H., Takahashi, N., Ohkawa, Y., Ji, S., Sugiyama, F., Fujio, K., Kumanogoh, A., Yamamoto, K., Kawasaki, N., Kurosaki, T., Takahashi, Y. and Furukawa, K. (2016) Sialylation converts arthritogenic IgG into inhibitors of collagen-induced arthritis. Nature Communications, 7, 11205.

Ossowski, S., Schneeberger, K., Clark, R.M., Lanz, C., Warthmann, N. et al. (2008) Sequencing of natural strains of *Arabidopsis thaliana* with short reads. Genome Research, 18, 2024–2033.

Pacurar, D.I., Pacurar, M.L., Street, N., Bussell, J.D., Pop, T.I., Gutierrez, L. and Bellini, C. (2012) A collection of INDEL markers for map-based cloning in seven Arabidopsis accessions. Journal of experimental botany, 63, 2491–2501.

Pidkowich, M.S., Klenz, J.E. and Haughn, G.W. (1999) The making of a flower: control of floral meristem identity in IT>Arabidopsis/IT>. Trends Plant Sci, 4, 64–70.

Pineiro, M. and Coupland, G. (1998) The control of flowering time and floral identity in Arabidopsis. Plant Physiol, 117, 1–8.

Schneeberger, K. (2014) Using next-generation sequencing to isolate mutant genes from forward genetic screens. Nat Rev Genet, 15, 662–676.

Schneeberger, K., Hagmann, J., Ossowski, S., Warthmann, N., Gessing, S. et al. (2009a) Simulataneous alignment of short reads against multiple genomes. Genome Biology, 10, R98.

Schneeberger, K., Ossowski, S., Lanz, C., Juul, T., Petersen A.H., Nielsen K.L., Jørgensen, J.E., Weigel D. and Andersen S.U. (2009b) SHOREmap: simultaneous mapping and mutation identification by deep sequencing. Nature Methods, 6, 550–1.

Shinya, T., Galis, I., Narisawa, T., Sasaki, M., Fukuda, H., Matsuoka, H., Saito, M. and Matsuoka, K. (2007) Comprehensive analysis of glucan elicitor-regulated gene expression in tobacco BY-2 cells reveals a novel MYB transcription factor involved in the regulation of phenylpropanoid metabolism. Plant Cell Physiol, 48, 1404–1413.

Siriwardana, N.S. and Lamb, R.S. (2012a) A conserved domain in the N-terminus is important for LEAFY dimerization and function in Arabidopsis thaliana. Plant J, 71, 736–749.

Siriwardana, N.S. and Lamb, R.S. (2012b) The poetry of reproduction: the role of LEAFY in Arabidopsis thaliana flower formation. Int J Dev Biol, 56, 207–221.

Sun, H.Q. and Schneeberger, K. (2015) SHOREmap v3.0: fast and accurate identification of causal mutaitons from forward genetic screens. Methods Mol. Biol., 1284, 381–95.

Weigel, D., Alvarez, J., Smyth, D.R., Yanofsky, M.F. and Meyerowitz, E.M. (1992) LEAFY controls floral meristem identity in Arabidopsis. Cell, 69, 843–859.

Winter, C.M., Austin, R.S., Blanvillain-Baufume, S., Reback, M.A., Monniaux, M., Wu, M.F., Sang, Y., Yamaguchi, A., Yamaguchi, N., Parker, J.E., Parcy, F., Jensen, S.T., Li, H. and Wagner, D. (2011) LEAFY target genes reveal floral regulatory logic, cis motifs, and a link to biotic stimulus response. Dev Cell, 20, 430–443.

Wu, X., Dinneny, J.R., Crawford, K.M., Rhee, Y., Citovsky, V., Zambryski, P.C. and Weigel, D. (2003) Modes of intercellular transcription factor movement in the Arabidopsis apex. Development, 130, 3735–3745.

