## Supplemental Figures and Tables for "The striking *flower-in-flower* phenotype of *Arabidopsis thaliana* Nossen (No-0) is caused by a novel *LEAFY* allele"

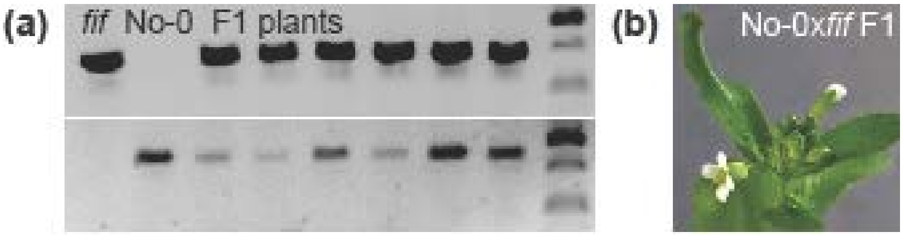


**Figure S1**. Proof of the successful backcross of the *fif* mutant with wild-type No-0.
**(a)** Genotyping PCR analysis of F1 plant material using a primer pair (sense: 5´-CGTGAAACTCAAGGCATTCTCTACTTC -3´; antisense: 5´-CGATTTCGACTTTAACCCGACCGG -3´) specifically amplifying the Ds transposon (upper row) and a primer pair (sense: see above; antisense: 5´-CGTACGTAGAACAACAGAGAATAAGC-3´) specifically amplifying the wild-type genomic region (lower row). **(b)** Representative image of the inflorescence and flower phenotype of F1 plants generated by the cross of the *fif* mutant with wild-type No-0 (♀No-0 x ♂*fif*). The reciprocal cross (♀*fif* x ♂No-0) provided the identical the results.

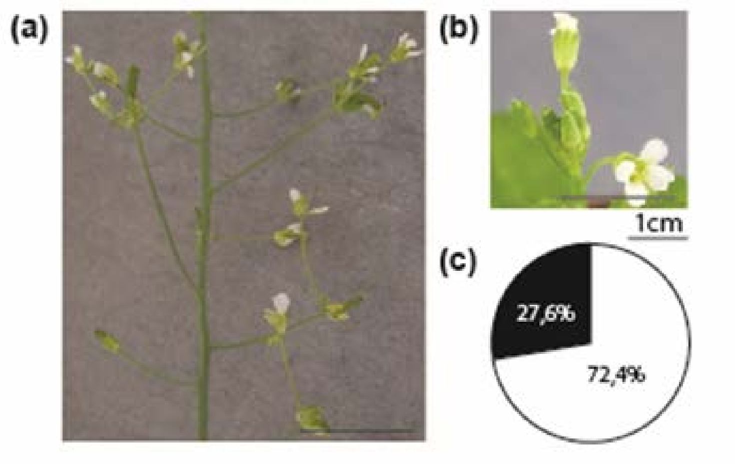


**Figure S2**. Distribution of the *fif* phenotype in the outcrossed F2 mapping population derived from plants of a F1 population generated by crosses of the *fif* mutant (No-0) with wild-type Col-0 (♀No-0 x ♂*fif*, ♀*fif* x ♂No-0). Representative images of the inflorescence of an outcrossed *fif* mutant **(a)** and wild-type individual **(b)**. **(c)** Distribution of plants showing either the *fif* mutant or wild-type floral phenotype within the outcrossed F2 mapping population. White circle outcut: plants with wild-type phenotype (72.4 %), black outcut: plants *fif* phenotype (27.6 %).Proof of the successful backcross of the *fif* mutant with wild-type No-0.

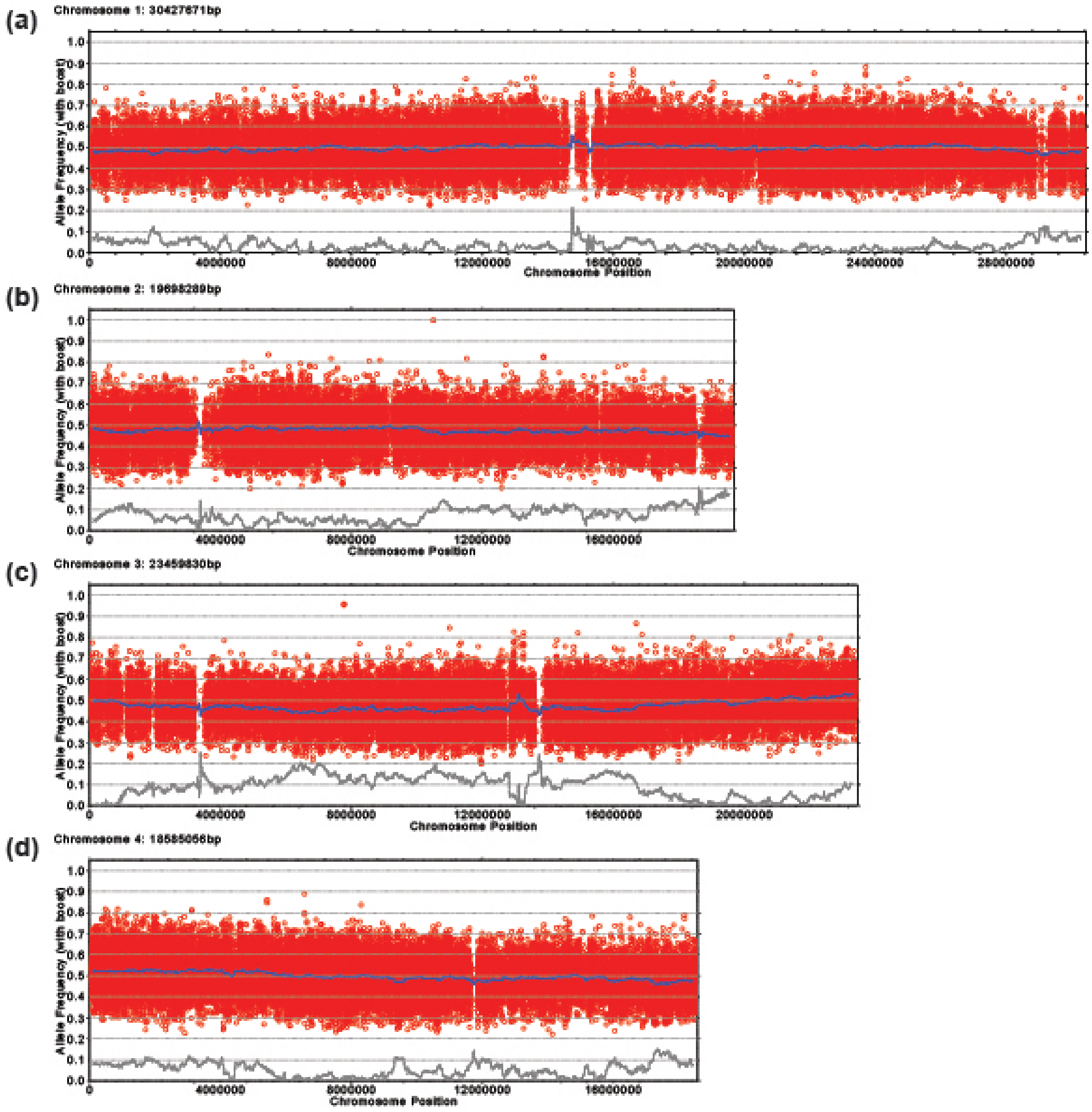


**Figure S3**. Allele frequency analysis of the Nos-0 allele within the recombinant mutant pool (unlinked chromosomes). Each red circle refers to a SNP marker distinguishing the Nos and Col genotypes. The blue line refers to a 200 kb sliding window analysis of the allele frequencies. The brown line would highlight potential mapping intervals (x-axis: genomic location; y-axis: Nos-0 allele frequency). **(a)** Chromosome 1. **(b)** Chromosome 2. **(c)** Chromosome 3. **(d)** Chromosome 4.

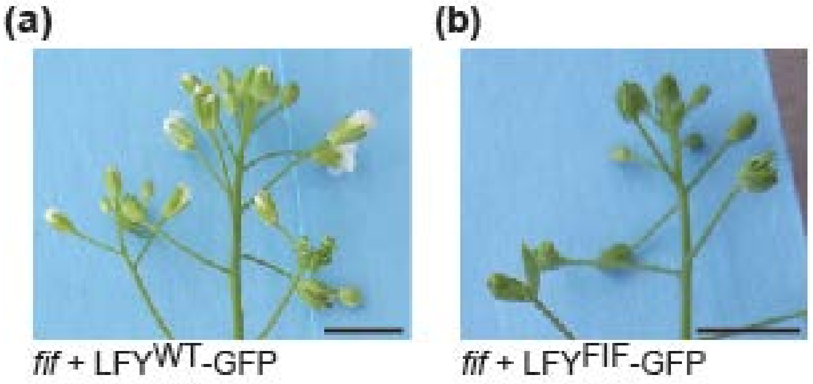


**Figure S4**. Complementation analysis of the *fif* mutant phenotype. *fif* mutant plants were transformed with binary constructs expressing, under the control of the UBQ10 promoter, either wild-type LFY-GFP or $LFY^{FIF}$-GFP. Whereas the expression of LFY-GFP almost fully complements the *fif* floral phenotype of the mutant **(a)**, no complementation was observed for the expression of $LFY^{FIF}$-GFP **(b)**.

| Name | | Sequence | Marker Type | Enzyme | bp *Col-0* | bp *No-0* |
|---|---|---|---|---|---|---|
| 7_1-8660 | S | GCGGCACAACCTAAATGAAA | INDEL | | 189 | 168 |
| | A | TGCATGCAATTATCACGTATG | | | | |
| 15_1-26627 | S | GCAATTCATCAGCAGGAGGT | INDEL | | 245 | 261 |
| | A | ATCAGGGAGCAAAATGCAAG | | | | |
| 20_2-12717 | S | AAAATGGGGCCTAATACGTTG | INDEL | | 403 | ~180 |
| | A | CAAAGGAAACACCTGCATCA | | | | |
| 4_3-3716 | S | TAATGGTGGCCCAATCTCAT | INDEL | | 1482 | 613 |
| | A | AATTCCAAATGGAGCCACAA | | | | |
| 16_3-20726 | S | GGGCCCATTTCAACTAAGGA | INDEL | | 149 | ~160 |
| | A | TCTCACAAGCCCAGTAAAAACT | | | | |
| 42_4-17544 | S | CACCATTGACATTTGATGCAC | INDEL | | 214 | 234 |
| | A | CCGTAGCTCCATTGGCTTAT | | | | |
| 1_5-1576 | S | CAGCTCCGACGATGATGATA | INDEL | | 363 | ~420 |
| | A | TGGAGTAATTGTTCCTTCACAAA | | | | |
| 37_5-22317 | S | GCATTGAAATAGTGTTTTTAACCAAA | INDEL | | 132 | 152 |
| | A | TGTTGGTTGCCACCTTATCA | | | | |
| 19_5-17388 | S | TTTTGCAAGTCCGTAGTCAATG | INDEL | | 110 | 121 |
| | A | TTTGGTTTTGGAATTTCTTTTG | | | | |
| 35_5-19138 | S | AACTCATGCAATGCGACATC | INDEL | | 182 | 164 |
| | A | CCCGTCCATGATCTGTTTCT | | | | |
| 37_5-22317 | S | GCATTGAAATAGTGTTTTTAACCAAA | INDEL | | 132 | 152 |
| | A | TGTTGGTTGCCACCTTATCA | | | | |
| S5-16 | S | CACGAGAGATACCTGCAAAACAG | dCAP | DraII | 160 | 134+26 |
| | A | CAAACGCTTTTGAAATCATGGGT**C**C | | | | |
| S5-24 | S | GTAATACACAACAATGGGGAG | dCAP | Esp3I | 244+43+26 | 287+26 |
| | A | CATATTCGAGTTCTGATGCACAC | | | | |
| 5-LFY | S | TATCTGTTCCACTTGTACGAA**GT**AT | dCAP | AccI | 150 | 129+21 |
| | A | CATAAATTTCAAGATAATGAACGGTC | | | | |
| 5-LFY#2 | S | TATCTGTTCCACTTGTACGAA**GC**AT | dCAP | SphI | 129+21 | 150 |
| | A | Same as 5-LFY | | | | |

**Table S1** Names and sequences of the INDEL and dCAP primers. Bold and underlined: introduced mismatches to incorporate an ecotype specific restriction site in the PCR product

| Name | | | sequence |
|---|---|---|---|
| pAP1 | S | Bio- | aaaaaGAAGGACCAGTGGTCCGTACaaaaa |
| | A | | tttttGTACGGACCACTGGTCCTTCttttt |
| pAP1m | S | Bio- | aaaaaGAAGGAAAAGTAATCCGTACaaaaa |
| | A | | tttttGTACGGATTACTTTTCCTTCttttt |
| C28M12 | S | Bio- | aaaaaaTTTATACTTGATCATaaCTTaaaa |
| | A | | ttttAAGttATGATCAAGTATAAAtttttt |

**Table S2** Sequences of the dsDNA oligonucleotides used in the DPI-ELISA.
